# No Semantic Information is Necessary to Evoke General Neural Signatures of Face Familiarity - Evidence from Cross-Experiment Classification

**DOI:** 10.1101/2022.06.08.495239

**Authors:** Alexia Dalski, Gyula Kovács, Géza Gergely Ambrus

## Abstract

Recent theories on the neural correlates of face identification stressed the importance of the available identity-specific semantic and affective information. However, whether such information is essential for the emergence of neural signal of familiarity has not yet been studied in detail.

Here, we explored the shared representation of face familiarity between perceptually and personally familiarized identities. We applied a cross-experiment multivariate pattern classification analysis (MVPA), to test if EEG patterns for passive viewing of personally familiar and unfamiliar faces are useful in decoding familiarity in a matching task where familiarity was attained thorough a short perceptual task. Importantly, no additional semantic, contextual, or affective information was provided for the familiarized identities during perceptual familiarization. Although the two datasets originate from different sets of participants who were engaged in two different tasks, familiarity was still decodable in the sorted, same identity matching trials.

This finding indicates that the visual processing of the faces of personally familiar and purely perceptually familiarized identities involve similar mechanisms, leading to cross-classifiable neural patterns.

## Introduction

In comparison to faces of unfamiliar people, personally relevant and highly familiar faces have been shown to be processed on a neural level in quantitatively and qualitatively dissimilar ways (Ramon and Gobbini 2018; Karimi-Rouzbahani et al. 2021). What exact factors shape these neuronal processes, and how they evolve as we get to know a person both in terms of sensory and social experience, is not fully understood (White and Burton 2022).

Several recent studies on the neural correlates of face-familiarity stressed the importance of the available identity-specific semantic and affective information. It has been consistently found that additional semantic information enhances performance in face recognition for short-term experimentally familiarized faces. For example, associating a novel face with additional information, such as the name or occupation of the person, enhances subsequent recognition performance (Schwartz and Yovel 2016). Furthermore, participants are better at remembering faces which they made conceptual decisions about, in contrast to faces with purely perceptual evaluations (Schwartz and Yovel 2018). Also, in a perceptual/semantic face-learning study by Kaufmann and colleagues (Kaufmann et al. 2009) participants remembered faces from short video clips better when simultaneously listening to short autobiographies. The ERP correlates of the semantic information were 700 ms following stimulus onset, suggesting the modulation of later, post-perceptual processing stages. With regard to the anatomical localization of semantic knowledge about people, the anterior temporal network and the hippocampus have been shown to be involved (Morton et al. 2021).

In a recent ERP study, Wiese and colleagues (2019) described a late, sustained familiarity effect (SFE) starting around 200 ms, and peaking between 400 and 600 ms following stimulus onset over occipito-temporal sites. In a subsequent study (Wiese et al. 2021) the authors investigated if this signal is modulated by the type and degree of familiarity. Their stimulus set contained images of the participant’s own face, personally familiar identities, liked, disliked and neutral celebrities, and unknown persons. They found that the SFE was present and similar in its characteristics for both personally familiar faces and for faces that were familiarized through media. As the SFE was modulated through the degree of familiarity, but not the mode of familiarization, the authors concluded that identity-specific semantic information plays an important role for the presence of a strong familiarity signal.

In our recent face-familiarization study (Ambrus et al. 2021) we set out to investigate the evolution of identity-related effects as previously unfamiliar faces become familiar. In this study three experiments were conducted: 1) a short perceptual exposure to the novel faces, 2) familiarization through media via watching a television series featuring the to-be-learned identities and 3) a personal familiarization experiment, requiring live, in-person interaction with the individuals on three occasions throughout one week. While we observed robust familiarity effects in the personal and media conditions, we found no clear differential pre- vs. post-familiarization identity-related changes.

This result is at odds with the results of Campbell and Tanaka (2021), who emphasized the importance of socially relevant, conceptual information in face learning. Their fast visual periodic stimulation study showed that personal familiarization spanning eight weeks with a previously unknown lab rotation partner, led to significant effects on the separation of familiar vs unfamiliar faces, predominantly in the occipitotemporal regions of the right hemisphere. The authors interpreted this finding as an identity-specific effect. They have argued that, in contrast to the short-term (one week long) personal familiarization period of Ambrus et al. (2021) a longer, socially more relevant familiarization phase is necessary to obtain enhanced identity representations of familiarized persons.

However, it is possible that although Campbell and Tanaka aimed to show identity-specific representations, their differential signal might in fact also be explained by familiarity effects, or the combination of signals arising from both the processing of face familiarity *and* identity. Whether a signal is related to a process characterized by identification “this is Bob’s face” (identity) or by recognition “this is a familiar face” (familiarity) is a challenging task without showing separable individuation-related signals in response to *multiple* newly learned identities. More importantly for our present purposes, one needs a control condition that involves no semantic, contextual, or affective, e.g., perceptual familiarization as the importance of these factors is otherwise hard to ascertain. To better understand the neural signals related to identity and familiarity processing, it is thus necessary to establish, what kind and amount of exposure to novel faces triggers the familiarity signals. Such an investigation, to our knowledge, has not yet been conducted.

Therefore, the current study aims at exploring whether even short term, purely perceptual face-identity learning can give rise to neural patterns that resemble those elicited by more extensive personal familiarization. Originally, perceptual learning did not lead to a robust familiarity signal, when running a within-subject classification procedure (Ambrus et al. 2021). Nevertheless, non-significant, chance-level decoding does not automatically rule out differential neural processing, as the quality of the results may strongly depend on, among other factors, the signal-to-noise ratio, classifier type and its hyperparameters, and the capacity of the sensors to pick up the relevant signals (Grootswagers et al. 2017).

In a subsequent report, we therefore reanalyzed the data of this study using a cross-experiment and cross-participant classification analysis on the three different familiarity conditions, with the aim of finding general neural signatures of face familiarity, irrespective of familiarization type, stimuli, and participants (Dalski et al. 2021). Such cross-experiment classification can potentially be more sensitive than within-subject classification methods, as it benefits from larger training datasets and is less confounded by idiosyncratic participant-level effects and stimulus properties. Iteratively training and testing classifiers on pairs of these datasets, we indeed found mutually cross-classifiable familiarity information starting around 200 ms post-stimulus onset, when training and testing across the media and personal familiarization experiments. However, in analyses involving perceptual familiarization, the results were more ambiguous: when the classifiers were trained on media or personal, and tested on the perceptual data, no shared familiarity signal was observed. Here it is important to note, that both the media and personal experiments involved a considerably longer, more context-rich familiarization phase, compared to the perceptual learning arm of the study.

This result thus may suggest that the general face-familiarity signature is modulated by the amount of biographical and semantic information that was only available in the media and personal, but not the perceptual experiment. However, as training on the perceptual and testing on the personal data also yielded a less pronounced, but significant results, the question of the necessity of social relevance, semantic, autobiographical, and affective information remains open. It is therefore important to establish whether these factors are indeed required for such shared, general face-familiarity effects to emerge.

To answer this question, neural patterns in response to faces learned through context-rich personal, and through context-poor, short-term perceptual familiarization, can be jointly examined. Here, we therefore made use of the personal familiarization dataset described in our previous reports (Ambrus et al. 2021; Dalski et al. 2021) and previously unreported data from the face-matching phase which concluded the perceptual face learning experiment (Ambrus et al. 2021). Here, we performed a cross-classification analysis by training on the personal familiarization and testing on the perceptual matching phase data, to test whether neural signals in this phase can be successfully categorized. In contrast to our previous analyses where data was recorded during passive exposure, this final face matching task required additional engagement with the faces presented, potentially leading to a stronger signal. Successful cross-classification in this case would confirm that a shared, general familiarity signal can arise even in the absence of semantic, social, and affective information about the familiarized identities.

## Methods

We conducted a cross-experiment classification analysis on two experiments reported in Ambrus et al. (2021). The experiments were conducted in accordance with the guidelines of the Declaration of Helsinki and were approved by the ethics committee of the Friedrich-Schiller-Universität Jena. Written, informed consent was obtained from all participants included in the studies. We trained classifiers on EEG ERP data recorded in the post-familiarization phase of the Personal Familiarization experiment and tested the classifiers’ performance on the previously unreported EEG data acquired during a two-alterative forced choice (2AFC) face matching task at the end of the Perceptual Familiarization experiment.

### Datasets

For details on stimulus presentation, data acquisition and preprocessing, see Ambrus et al. (2021) and Dalski et al. (2021). **Figure 1**. Shows the experimental design for the parts of the two experiments from which data was used in the present analysis. In short, all stimuli were color, ambient face images, depicting initially unfamiliar identities. The images were eye-aligned, cropped to center on the inner features of the face, and were presented centrally on a uniform gray background in a pseudorandom order. During the decoding phases of the experiments, the volunteers were given a simple target detection task (button press at the detection of slightly rotated images) to ensure maintained attention throughout the recording. No participant took part in both experiments.

**Figure 1.**
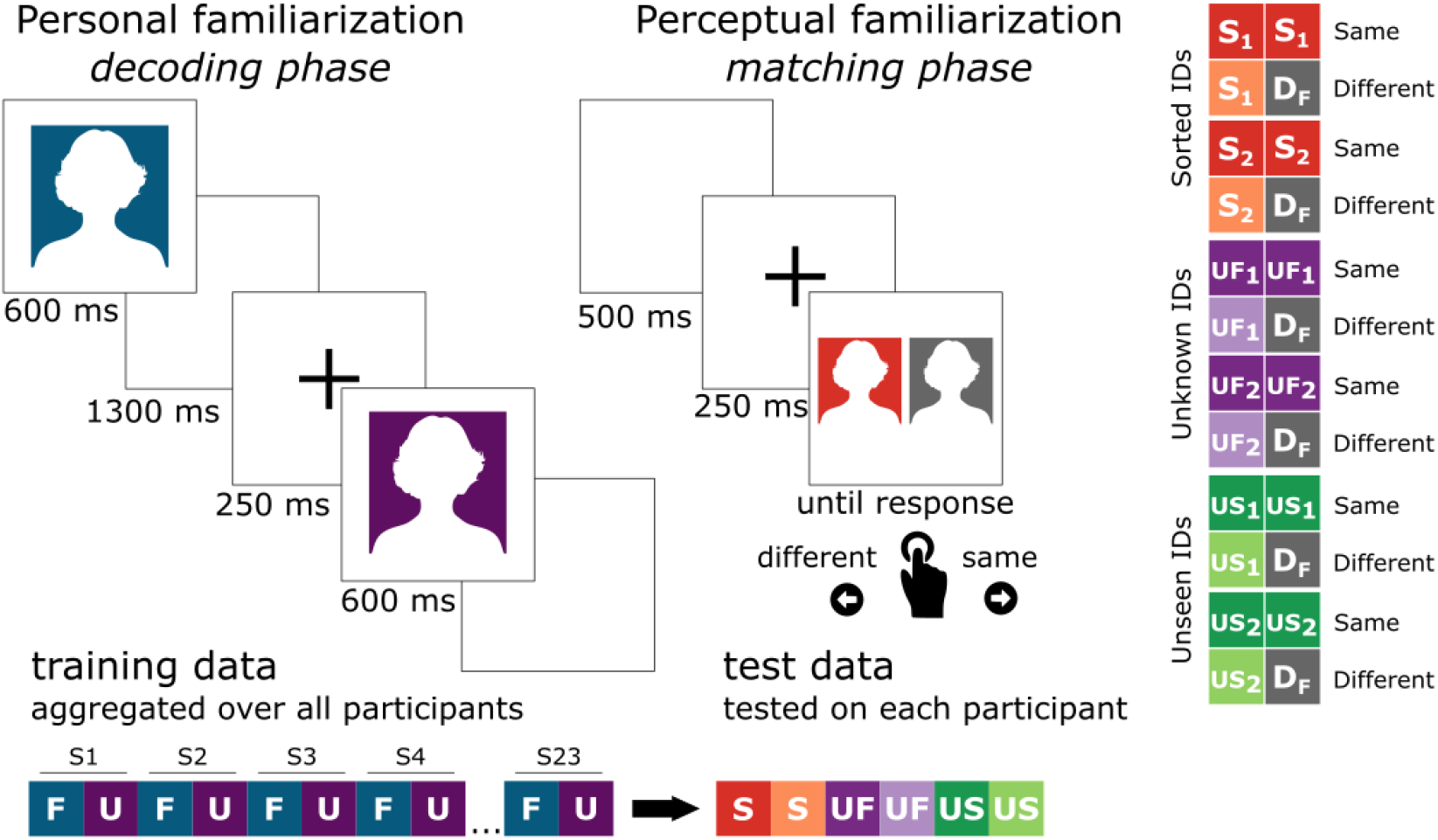
Experimental design. *Left:* incidental exposure trials in the Personal Familiarization experiment. In these trials one image of a personally familiarized (F) or an unfamiliar (U) person was shown on the screen for 600 ms. EEG data aggregated over all participants from this experiment served as the training set. *Right:* the matching task at the end of the Perceptual Familiarization experiment that provided the test dataset for cross-classification. Here, two images of either the same identity, or two different identities were shown simultaneously. These could be either familiarized (S_1_, S_2_), unfamiliar (but seen during the previous decoding phase, UF_1_, UF_2_) or entirely new, previously unseen identities (US_1_, US_2_). In the ‘different’ trials an image of a trial-unique Foil identity (D_F_) was used. The stimuli were present on screen until a same/different response was given using the arrow keys. See **Figure 2** for the stimuli in the matching task.

Below is a short overview of the experiments that provided the training and testing datasets:

#### Personal Familiarization

The dataset includes data from 23 participants. Stimuli were four previously unknown female identities, with 10 ambient images each. During the pre- and post-familiarization EEG recording phases, one image was shown 22 times (see **Figure 1**, left). The familiarization phase consisted of ca. one-hour personal meetings with two of the four identities on 3 consecutive days between pre- and post-familiarization EEG recordings. The to-be-familiarized identities were the same for all participants.

#### Perceptual Familiarization

The dataset includes data from 42 volunteers. This experiment consisted of three phases, each of which involved a new set of photographs of the persons presented. The *sorting phase* consisted of 120 trials, two repetitions of the 30 – 30 images of two to-be-familiarized identities. The images were presented sequentially. The task of the participant was to sort these images into two identities using the left and right arrow keys. Other than the number of identities the participants needed to sort, we provided no further information (such as name, occupation, etc.) about the stimuli. The familiarization phase lasted 5.29 minutes on average (SD=1.29)

During the *decoding phase* in a total of 1600 trials, 10 – 10 novel photographs of these familiarized identities were presented, along with 10 – 10 images of two unfamiliar identities, randomly intermixed. The stimulus presentation time was 600 ms. This phase lasted 94.51 minutes on average (SD=7.68).

Finally, in each of the 120 trials of the *matching phase* (**Figure 1**, right), participants were asked to perform a two-alternative forced choice matching task of two face images, where a new set of photographs of the familiarized and unfamiliar images were presented side-by-side horizontally, depicting the same identity, or another, trial-unique, unrelated person (foil). Furthermore, the participants were tested on two additional, heretofore unseen identities the same way. In summary, each identity (2 familiarized, 2 unfamiliar, 2 unseen) was presented in 10 same-identity and 10 different-identity trials. The images remained on screen until a response was given. The left-right position of the two images was randomly chosen in each trial. At the beginning of the experiment, the participants practiced the face matching task on 10 trials, in which the stimuli consisted of images of famous male identities. Feedback was given during practice, but not during the actual experiment. This matching phase lasted 8.13 minutes on average (SD=3.60). Data from this phase of the study has not been reported previously.

The core stimulus set consisted of face-images of four previously unknown female identities. For each participant, the two to-be-familiarized identities were randomly chosen from the 6 possible permutations of these four identities. The two additional unseen identities in the matching task were the identical for all participants. For use as foils in the different identity trials, 60 photographs of 60 additional, different persons were used. All identities were female and chosen to be of similar general appearance (e.g., similar age and hair color, see **Figure 2**).

**Figure 2.**
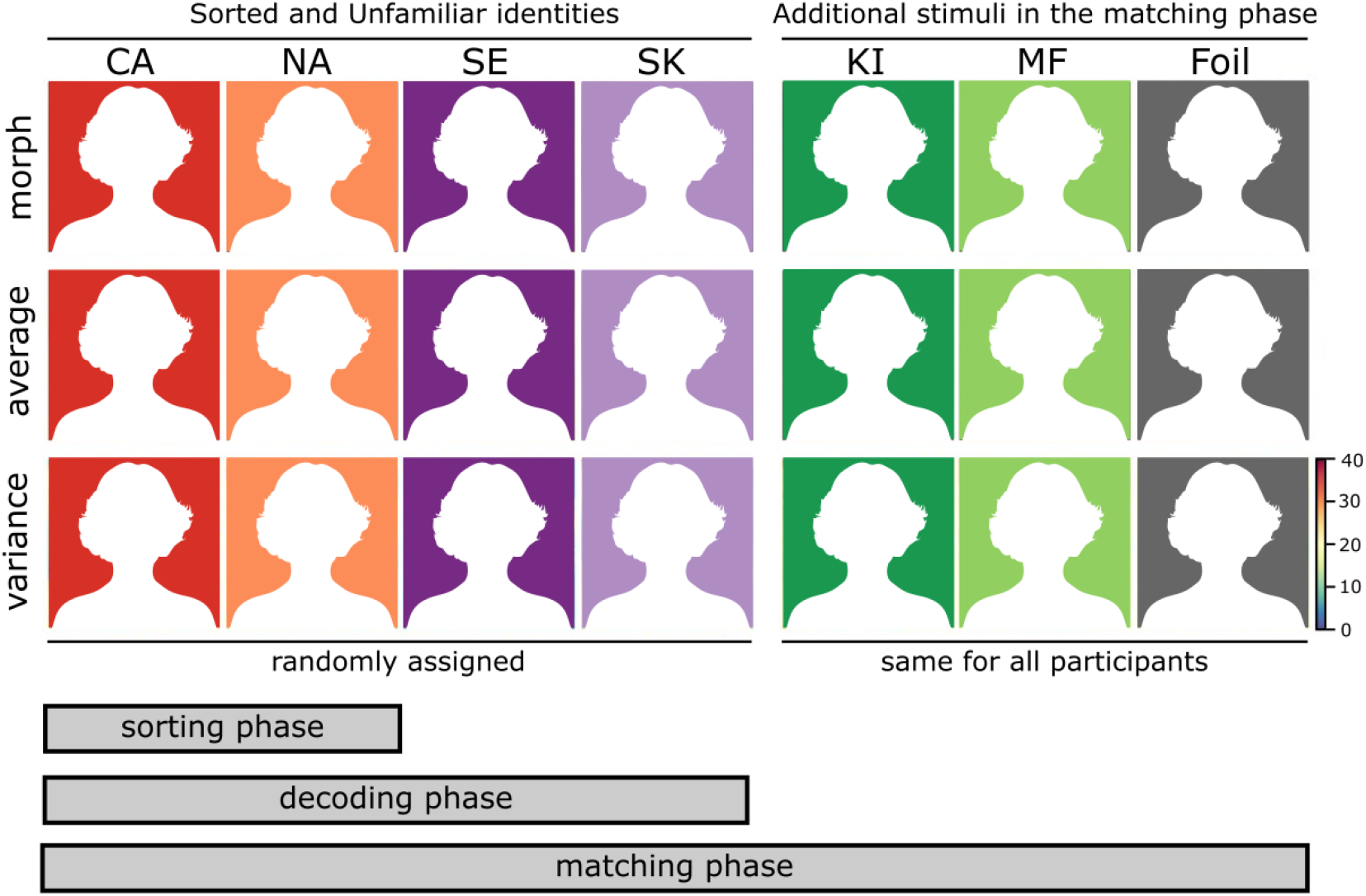
Stimuli in the Perceptual familiarization experiment. Top panel: Morphs of the face-images of the identities in the matching task stimulus set (Sorted and Unfamiliar: CA: Adél Csobot, NA: Anikó Nádai, SK: Kata Sarka, SE: Erika Szabó; Unseen: KI: Katarina Ivanovska, MF: Fruzsina Maranec). The Foil image is a morph composite of all images used as different identities in the matching task. Middle panel: Pixel-wise averages of the images. Lower panel: Variance across the images, with warmer colors indicating higher pixel-wise variance. Images of two identities, randomly chosen for each participant, served as the to-be-familiarized (Sorting) stimuli. Novel images of these two identities, together with the remaining two (Unfamiliar) identities, were shown in the decoding phase of the session. Finally, novel (i.e., not before presented) images of the Sorted and Unknown identities, together with two additional, previously Unseen identities, formed the stimulus set in the matching phase.

### Analysis

#### Face sorting and matching performance

Improvement in face-sorting was evaluated by comparing the consistency of responses to the images of the two Sorting identities in the first and the last 20 trials in the sorting phase of the Perceptual familiarization experiment. A two-tailed paired-sample t-test was used. Matching performance and reaction times were submitted to repeated measures ANOVAs with Stimulus Familiarity (Familiarized/Sorted; Unfamiliar; Unseen) and Match Condition (Same; Different) as within-subject factors.

#### Cross-classification analysis

EEG was recorded using a 64-channel Biosemi Active II system (512 Hz sampling rate) in a dimly lit, electrically shielded, and sound–attenuated chamber. Data were bandpass filtered between 0.1 and 40 Hz, segmented between −200 and 1300 ms and baseline corrected to the 200 ms preceding the stimulus presentation. The data were downsampled to 100 Hz, with no artifact rejection performed (Grootswagers et al. 2017). Processing was carried out using MNE-Python (Gramfort et al. 2013).

The analysis pipeline was based on Dalski et al. (2021). For training, we used to post-familiarization phase of the Personal Familiarization experiment reported in Ambrus et al. (2021). The test dataset was taken from the final face-matching test of the Perceptual Familiarization experiment described in the same report.

Epoched EEG data from all participants in the post-familiarization phase of the Personal Familiarity experiment, from all trials and all channels, was concatenated. Trials with the presentation of the two familiarized and the two unfamiliar identities were re-coded to ‘familiar’ and ‘unfamiliar’ as labels used for classification. At each of the 150 time points (−200 to 1300 ms relative to stimulus presentation onset, sampled at 100 Hz), linear discriminant analysis classifiers (LDA) were trained to categorize familiarized and unfamiliar identities. These classifiers were then used to calculate prediction performance (classified as ‘familiar’) in the matching phase of the Perceptual Familiarity experiment, for each participant separately.

Trials in the Perceptual Matching phase were divided into six categories along the Familiarity (Sorted, Unfamiliar, Unseen) and Match Condition (Same, Different) factors. For each of these categories, a separate cross-classification profile was calculated based on the ratio of ‘familiar’ classification. Several measures were taken to reduce noise in this dataset. Only correct responses were entered into the cross-classification analyses. To further reduce the noise due to volunteers with a small number of correct trials, in each category, only data from participants with 10 or more correct trials were analyzed (Sorted Same: *n*=38; Sorted Different: *n*=42; Unfamiliar Same: *n*=38; Unfamiliar Different: *n*=41; Unseen Same: *n*=38; Unseen Different: *n*=42). A 30 ms moving average (3 consecutive time-points) was used on all cross-experiment classifier performance data at the participant level in order to increase signal-to-noise ratio (Kaiser et al. 2016; Ambrus et al. 2019). Results of the time-resolved analyses were tested for statistical significance using two-tailed cluster permutation tests.

##### Spatio-temporal searchlight

First, we conducted a searchlight analysis, in which all channels were systematically tested separately by training and testing on data originating from the given sensor and adjacent electrodes. For each channel and its neighbors, a time-resolved analysis was conducted. Two-tailed spatio-temporal cluster permutation tests against chance level (50%), with 10,000 iterations, were used for the purposes of statistical inference.

##### Regions of interest analysis

Data form all sensors, and pre-defined regions of interest, were subjected to time-resolved cross-classification. To construct the ROIs, similarly to Ambrus et al. (2019, 2021), and Dalski et al. (2021), we defined six scalp locations along the medial (left and right) and coronal (anterior, center, and posterior) planes. We used the subsets of channels in these regions for training and testing in separate analyses. Cross-classification performance was evaluated using two-tailed, one-sample cluster permutation tests (10,000 iterations) against chance (50%).

## Results

### Behavioral results

#### Sorting performance

Participants in our Perceptual familiarization experiment were familiarized through a short, 120 trial card sorting task at the beginning of the session. Performance in this task improved by 20% from 65±11% in the first to 85±14% in the last 20 trials (*t*=9.176, *p*<0.001, Cohen’s *d*=1.416), indicating that the participants’ performance to tell the two identities apart improved throughout the task.

#### Matching performance

The effect of perceptual familiarization with the two Sorted identities was evaluated against that of the two Unfamiliar and two previously Unseen identities in a face matching task at the end of the experiment.

The repeated measures ANOVA performed on the accuracy data yielded a main effect of Stimulus Familiarity: *F*_2,82_ = 8.275, *p* < 0.001, η^2^p = 0.168. Performance was highest for the Sorted identities (mean: 0.79, ±SD: 0.10) and differed significantly from the Unfamiliar, but not from the unseen identities. A main effect of Match Condition was also observed, *F*_1,41_ = 20.153, *p* < 0.001, η^2^p = 0.330, with a higher accuracy for Different compared to Same identity trials. The interaction between Stimulus Familiarity and Match Condition was not shown to be statistically significant *F*_2,82_ = 2.804, *p* = 0.066, η^2^p = 0.064.

The same analysis on reaction times also yielded a significant main effect of Stimulus Familiarity: *F*_2,82_ = 4.186, *p* = 0.019, η^2^p = 0.093. Reaction times were significantly shorter for the Sorted compared to the Unfamiliar identities. Reaction times for previously Unseen identities were not statistically different from that observed for Sorted identities but differed from that in the Unfamiliar condition. A main effect of Match Condition was also observed, *F*_1,41_ = 5.347, *p* = 0.026, η^2^p = 0.115, with slower responses for Same as opposed to Different identity trials. Finally, the interaction between Stimulus Familiarity and Match Condition was shown to be statistically significant *F*_2,82_ = 5.347, *p* = 0.007, η^2^p = 0.115.

Summaries of the accuracies and reaction times in the different matching conditions in all participants are reported in **Table 1**.

**Table 1.** Results of the face matching task in the Perceptual Familiarization experiment. (all participants, *n* = 42). A) Descriptive statistics of matching accuracies and reaction times. B) Results of paired samples *t*-tests comparing same and different identity conditions between and across the three familiarity types (Sorted IDs, Unfamiliar IDs and previously Unseen IDs)

| Accuracy (%) |  |  |  |  |  | Reaction time (s) |  |  |  |  |
| --- | --- | --- | --- | --- | --- | --- | --- | --- | --- | --- |
|  | mean | sd | median | minimum | maximum | mean | sd | median | min | max |
| Sorted ID, Same | 75.357 | 16.616 | 75 | 25 | 95 | 3.424 | 1.939 | 2.887 | 1.448 | 11.71 |
| Sorted ID, Different | 82.976 | 11.265 | 85 | 55 | 100 | 3.44 | 1.911 | 2.937 | 1.417 | 9.812 |
| Unfamiliar ID, Same | 65.833 | 14.267 | 65 | 30 | 95 | 3.935 | 2.194 | 3.293 | 1.46 | 11.586 |
| Unfamiliar ID, Different | 80.119 | 13.99 | 80 | 45 | 100 | 3.429 | 1.75 | 2.804 | 1.471 | 8.948 |
| Unseen ID, Same | 69.762 | 15.616 | 75 | 40 | 95 | 3.576 | 1.737 | 3.139 | 1.479 | 8.907 |
| Unseen ID, Different | 83.214 | 11.833 | 85 | 55 | 100 | 3.395 | 1.817 | 2.997 | 1.371 | 10.681 |
|  | mean | sd | median | min | max | mean | sd | median | min | max |
| Same | 70.3 | 12.2 | 72.5 | 43.3 | 91.7 | 3.645 | 1.905 | 3.194 | 1.51 | 10.734 |
| Different | 82.1 | 10.8 | 83.3 | 55 | 1 | 3.421 | 1.777 | 2.837 | 1.441 | 9.714 |
|  | mean | sd | median | min | max | mean | sd | median | min | max |
| Sorting | 79.167 | 10.071 | 77.5 | 52.5 | 95 | 3.432 | 1.869 | 3.02 | 1.476 | 10.761 |
| Unfamiliar | 72.976 | 8.856 | 72.5 | 50 | 87.5 | 3.682 | 1.949 | 3.195 | 1.525 | 9.997 |
| Unseen | 76.488 | 9.863 | 77.5 | 47.5 | 95 | 3.486 | 1.717 | 3.029 | 1.425 | 9.608 |

| Accuracy |  |  |  |  | Reaction time |  |  |  |
| --- | --- | --- | --- | --- | --- | --- | --- | --- |
|  | t | p | Cohen's d | difference | t | p | Cohen's d | difference |
| Sorted ID, Same – |  |  |  |  |  |  |  |  |
| Sorted ID, Different | -2.468 | 0.018* | -0.381 | -7.619±20.007 | -0.111 | 0.912 | -0.017 | -0.016±0.927 |
| Unfamiliar ID, Same – |  |  |  |  |  |  |  |  |
| Unfamiliar ID, Different | -4.205 | <0.001*** | -0.649 | -14.286±22.019 | 4.381 | <0.001*** | 0.676 | 0.506±0.749 |
| Unseen ID, Same – |  |  |  |  |  |  |  |  |
| Unseen ID, Different | -4.48 | <0.001*** | -0.691 | -13.452±19.459 | 1.272 | 0.211 | 0.196 | 0.181±0.922 |
| Sorted ID, Same – |  |  |  |  |  |  |  |  |
| Unfamiliar ID, Same | 3.617 | <0.001*** | 0.558 | 9.524±17.065 | -3.701 | <0.001*** | -0.571 | -0.511±0.895 |
| Sorted ID, Same – |  |  |  |  |  |  |  |  |
| Unseen ID, Same | 2.043 | 0.047* | 0.315 | 5.595±17.745 | -1.169 | 0.249 | -0.18 | -0.152±0.844 |
| Unfamiliar ID, Same – |  |  |  |  |  |  |  |  |
| Unseen ID, Same | -1.684 | 0.1 | -0.26 | -3.929±15.123 | 3.027 | 0.004** | 0.467 | 0.359±0.768 |
| Sorted ID, Different – |  |  |  |  |  |  |  |  |
| Unfamiliar ID, Different | 1.974 | 0.055 | 0.305 | 2.857±9.38 | 0.098 | 0.922 | 0.015 | 0.011±0.729 |
| Sorted ID, Different – |  |  |  |  |  |  |  |  |
| Unseen ID, Different | -0.144 | 0.886 | -0.022 | -0.238±10.704 | 0.396 | 0.694 | 0.061 | 0.045±0.731 |
| Unfamiliar ID, Different – |  |  |  |  |  |  |  |  |
| Unseen ID, Different | -1.68 | 0.101 | -0.259 | -3.095±11.943 | 0.293 | 0.771 | 0.045 | 0.034±0.744 |
|  | t | p | Cohen's d | difference | t | p | Cohen's d | difference |
| Same - Different | -4.489 | <0.001*** | -0.693 | -11.786±17.014 | 2.312 | 0.026* | 0.357 | 0.224±0.627 |
|  | t | p | Cohen's d | difference | t | p | Cohen's d | difference |
| Sorted ID - Unfamiliar ID | 4.557 | <0.001*** | 0.703 | 6.19±8.804 | -2.471 | 0.018* | -0.381 | -0.25±0.655 |
| Sorted ID - Unseen ID | 1.616 | 0.114 | 0.249 | 2.679±10.741 | -0.602 | 0.55 | -0.093 | -0.054±0.579 |
| Unfamiliar ID - Unseen ID | -2.269 | 0.029* | -0.35 | -3.512±10.031 | 2.414 | 0.02* | 0.372 | 0.196±0.527 |

### Searchlight analysis

Results of the searchlight analysis are reported in **Figure 3**. Only the Sorted, Same Identity condition yielded a significant spatio-temporal cluster in the searchlight analysis (cluster *p* = 0.0003, peak time at 560 ms over PO9, peak *t* = 5.906, peak Cohen’s *d* = 0.958). This effect included a temporal cluster starting at 170 ms following stimulus onset and lasted until the end of the epoch. The effect was most prominent over occipito-temporal and anterior sensors, mainly in the left hemisphere.

**Figure 3.**
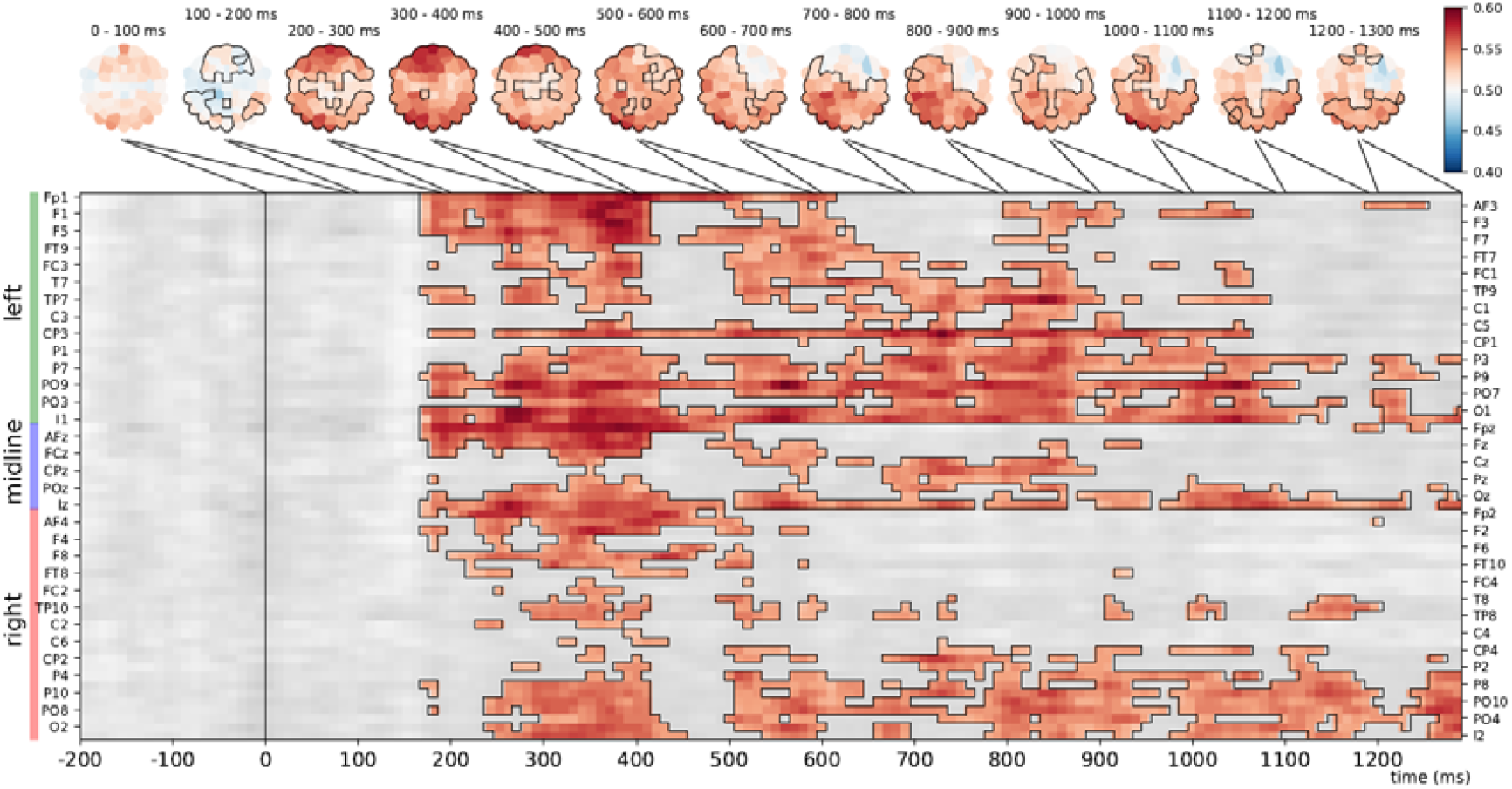
Searchlight analysis in the Sorted, Same Identity condition. *Top*. Scalp-maps showing mean classifier performance, averaged for each channel in successive 100 ms time-bins. Outlines indicate channels that were part of the significant spatio-temporal cluster within the bin. *Bottom*. The extent of the significant spatio-temporal cluster across channels and time-points. Two-tailed spatio-temporal cluster permutation test, cluster *p* = 0.0003

### Regions of Interest analysis

Results of the time-resolved ROI analyses are presented in **Figure 4**, see **Table 2** for detailed statistics. Time-resolved ROI analyses have revealed significant cross-experiment classification performance over all electrodes, and all six pre-defined ROIs in the Sorted, Same Identity condition. Short, relatively early significant clusters were seen in the left hemisphere in the anterior (200 to 320 ms) and posterior (170 to 260 ms) regions in the case of Sorted, Different identity condition. No significant effects were observed for the other conditions.

**Figure 4.**
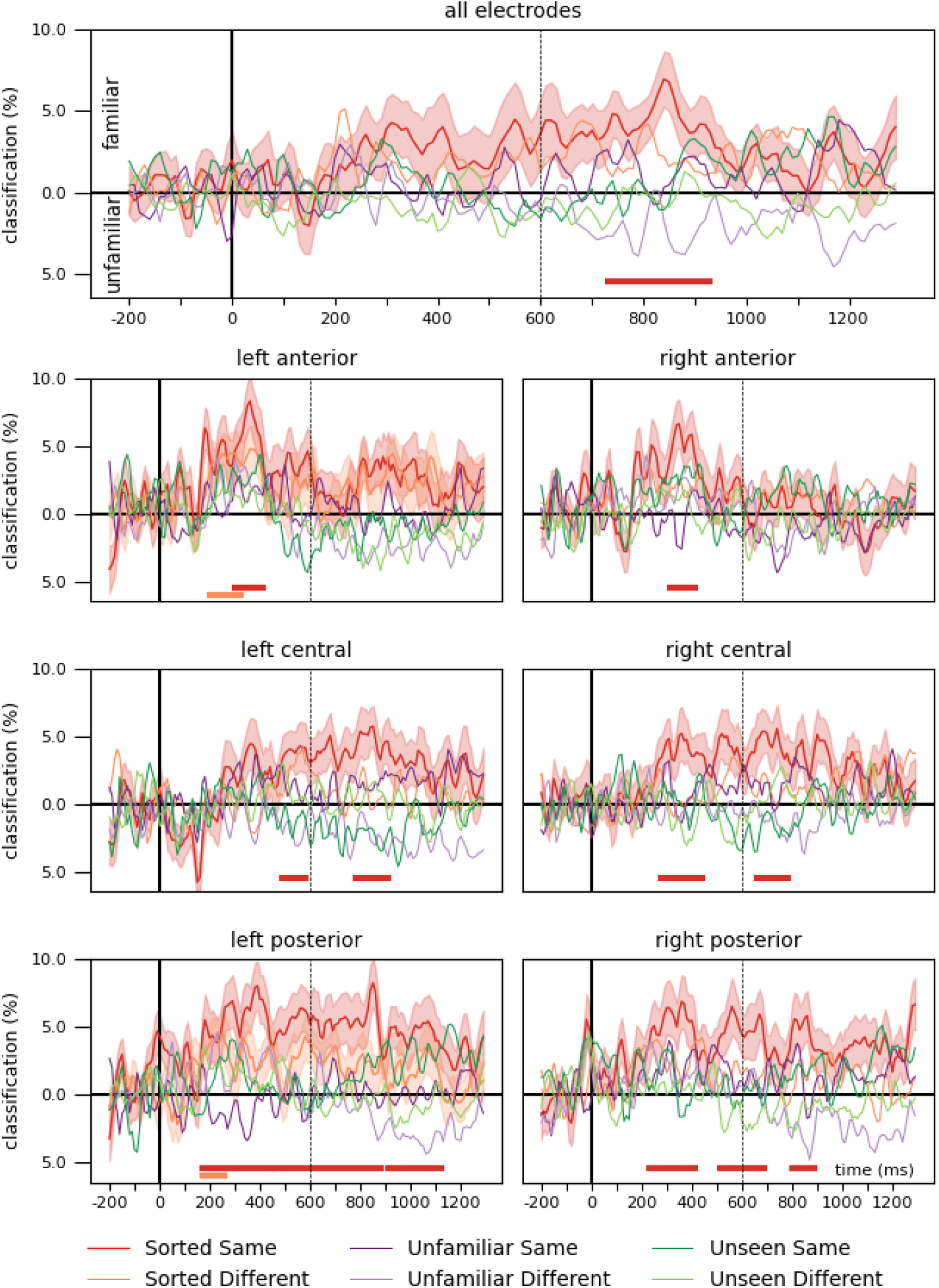
Time-resolved cross-classification performance on all electrodes and pre-defined regions of interest. Classifier performance when trained on ERPs recorded during passive viewing of personally familiar and unfamiliar faces in the Personal Familiarization experiment and tested on the ERPs in the face-matching task of the Perceptual familiarization study. Horizontal markers denote clusters with significantly different decoding accuracies against chance (two-tailed cluster permutation tests, *p* < 0.05). The vertical dashed line at 600 ms denotes stimulus offset in the training data. Detailed statistics can be found in **Table 2**.

**Table 2.**
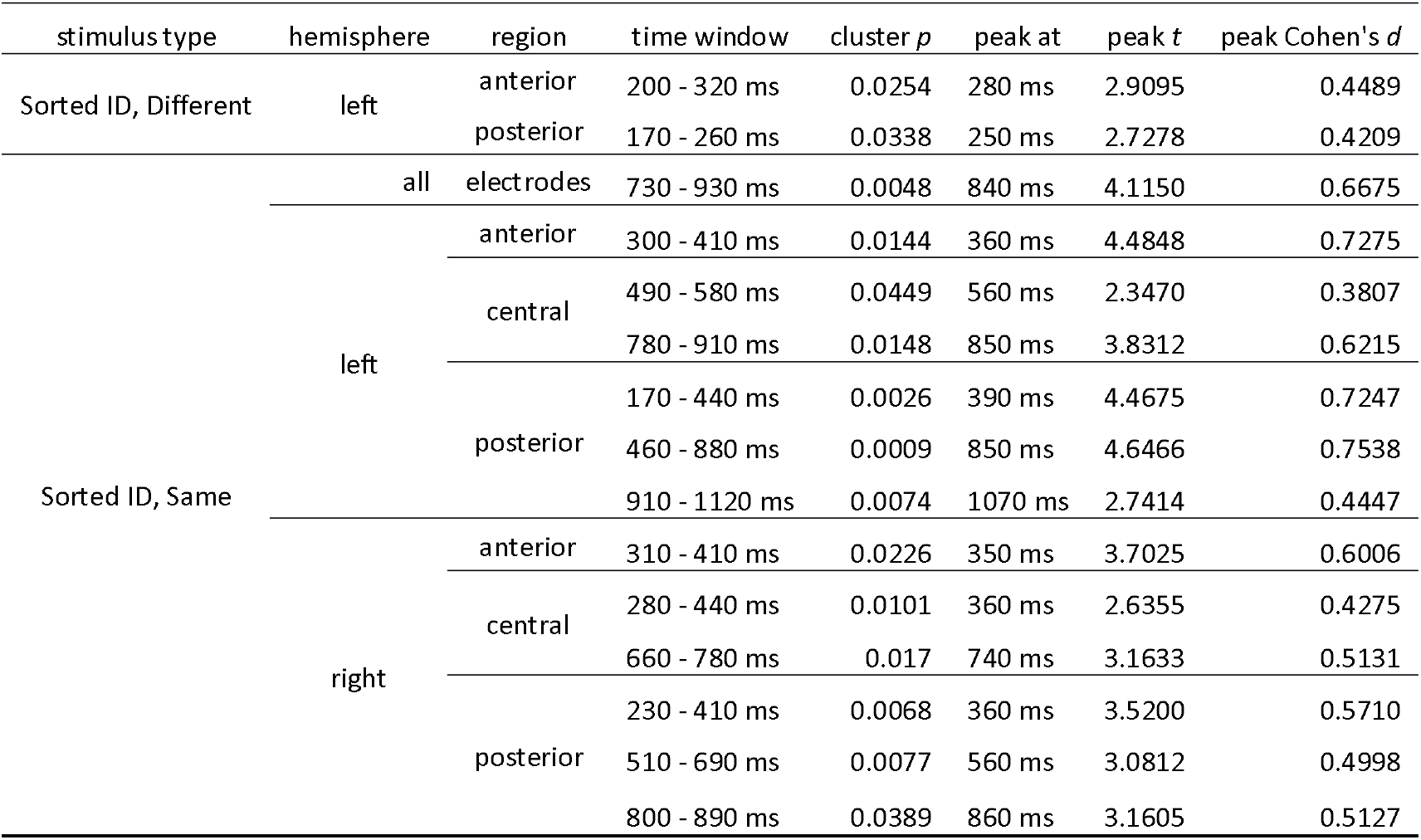
Results of the time-resolved cross-classification performance on all electrodes and pre-defined regions of interest. Clusters flagged as significant, two-tailed cluster permutation tests against chance.

Although the familiarity cross-classification performance in the Sorted, Same Identity condition in all electrodes was consistently over the chance level, the cluster permutation test flagged only a late temporal window, between 730 and 930 ms, as significant. Earlier time windows were indicated in various regions of interest, especially in the anterior electrodes, where two short clusters were observed bilaterally (left: 300 to 410 ms, right: 310 to 410 ms). Central sites each yielded two clusters (left: 490 to 580 ms and 780 to 910 ms, right: 280 to 440 ms, and 660 to 780 ms). Interestingly, a more robust effect was seen in left posterior ROI, which yielded three almost contiguous significant clusters between 170 and 1120 ms (170 to 440 ms; 460 to 880 ms; 910 to 1120 ms), while the three significant clusters in the right posterior region appeared later and were more restricted in time (230 to 410 ms; 510 to 690 ms; 800 to 890 ms).

To better visualize the trends in the cross-classification time-course, for each trial, we averaged classifier performance in each participant in the post-stimulus-onset time range, then aggregated these for each condition (**Figure 5**). As stimulus duration and the response window was not limited in the matching task, we reasoned that a combined measure of the magnitude and duration of the signal, i.e., averaging across time, is a useful metric to observe trends in classification performance. The results of this procedure largely mirrored those in the time-resolved analyses. Sorted, Same identity trials were classified as familiar significantly above chance level in all regions of interest, except over the right anterior sensors. In addition, the Sorted, Different identity stimuli were classified as familiar significantly above chance in the left posterior ROI, and Unfamiliar, Different Identity trials as unfamiliar in the left central region of interest. No other significant difference from chance was observed.

**Figure 5.**
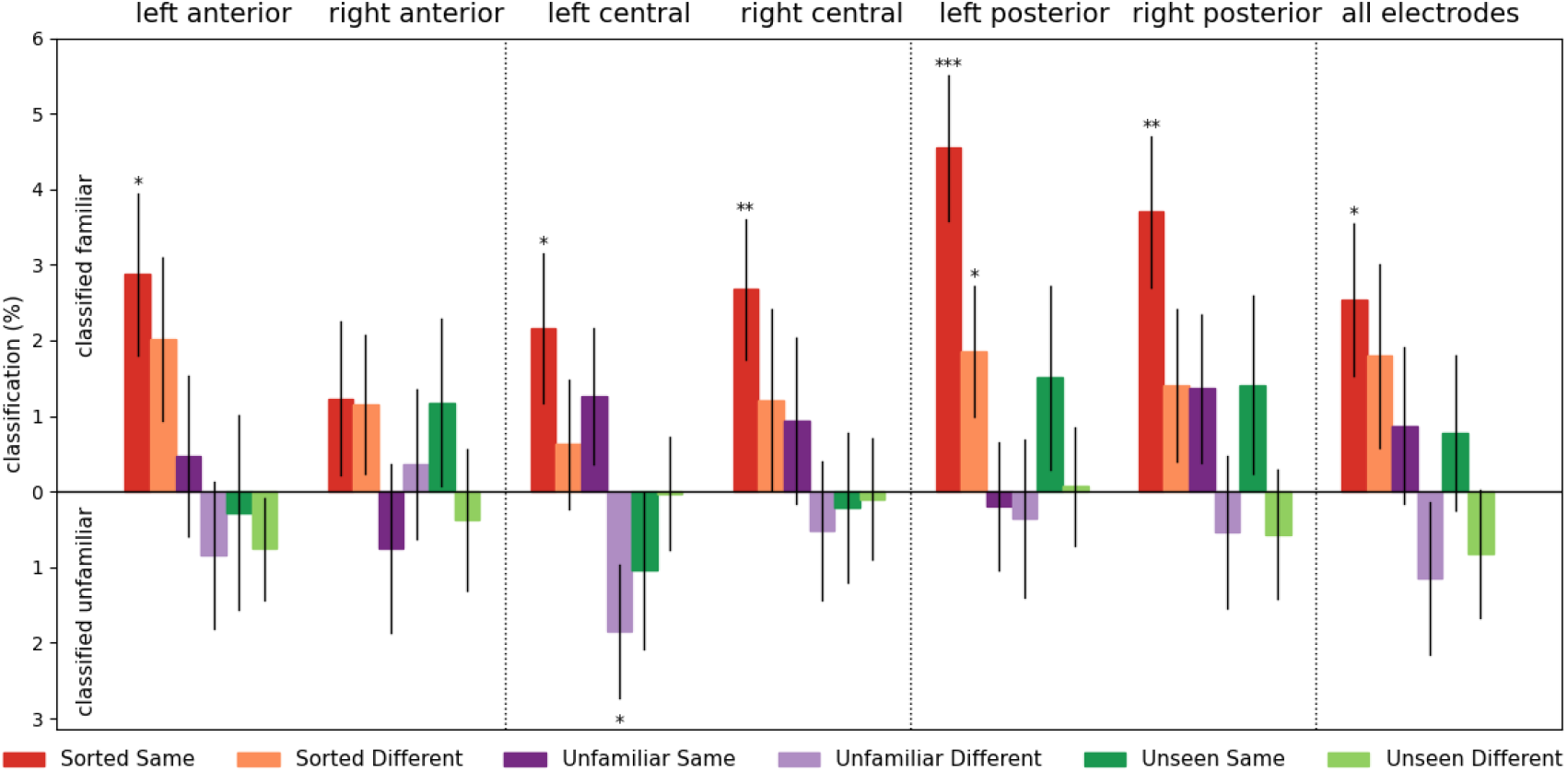
Classifier cross-classification performance, averaged across time. For each participant in each trial, classification performance was averaged from stimulus onset to the end of the epoch, then averaged for stimulus types and aggregated across participants. Error bars denote standard errors. One-sample two-tailed *t*-test against chance, \**p*_uncorrected_ < 0.05, \*\**p*_uncorrected_ < 0.01, \*\*\**p*_uncorrected_ < 0.001

## Discussion

Our study sought to test if a general neural signature of face-familiarity can emerge for faces that were learned without any semantic information. For this purpose, we tested if the strong familiarity signals observed for passive viewing of personally familiar faces in the EEG patterns generalize to purely perceptually familiarized faces in a face-matching task. The major findings of this study are the following: 1) The behavioral results in the face sorting and matching tasks imply that participants can learn image-independent representations through a short perceptual learning task; 2) successful cross-dataset classification for face familiarity is possible between experiments that use different tasks. 3) In the face matching task, trials with both stimuli depicting the same, familiarized identity, led to a robust cross-classifiable familiarity signal.

### Matching performance for familiarized identities implies the formation of image-independent representations

Performance in face matching indicates that a short sorting task was sufficient to enhance matching performance and lower reaction times for Sorted, when compared to Unfamiliar identity trials. This effect was observable one and a half hours after the completion of the familiarization task, and despite the incidental, repeated exposure to the 10 – 10 images of the two Unfamiliar identities. This indicates that our short, perceptual learning task was sufficient to tell the Sorted identities apart, and crucially, tell images of them together, implying the formation of image-independent identity representations (Burton et al. 2011; Andrews et al. 2015; Ambrus et al. 2017; Young and Burton 2017). On the other hand, passive viewing of images of the Unfamiliar identities did not lead to such effects.

### Potential effect of task on the decodability of neural patterns

Despite a clear learning effect in the face matching task, in our original study (Ambrus et al. 2021) we found little indication of a familiarity signal in a within-participant representational similarity analysis during the decoding phase. On the other hand, in our subsequent cross-experiment analysis (Dalski et al. 2021), classifiers trained on data in the Perceptual Familiarization experiment successfully predicted face familiarity in the Personal Familiarization experiment, suggesting that familiarity information is already present in the EEG. No such robust effect was seen in the other direction, i.e., training on Personal and testing on Perceptual data.

In our present analysis, in contrast to the decoding phase of the Perceptual Familiarization experiment where no identity-related task had to be performed by the participants, the matching task involved active engagement with the stimuli that could have facilitated familiarity-related processing of the novel images of the sorted identities. Similar task-dependent modulation of decodability has been observed in several studies previously. In an fMRI speaker/vowel classification test, task dependency modulated the informative neural response patterns, reflecting the top-down enhancement of relevant sound representations (Bonte et al. 2014). Yip et al. (2022) found that representations of the same objects led to quantitatively and qualitatively different EEG decoding time-courses across different task contexts, while another study (Hubbard et al. 2019) found that using a cued task-switching paradigm, information regarding relevant representations can be decoded from EEG. In an identity/facial expression categorization task, explicit processing of identity led to enhanced decodability of that dimension (Smith and Smith 2019). We argue that in our matching task, the relevance of image-independent identity matching might have amplified familiarity-related signals that could be successfully decoded using classifiers trained on the Personal Familiarization experiment data.

### Cross-experiment classification is feasible between different tasks

Multivariate cross-classification is increasingly used to investigate similarity among neural patterns recorded in different cognitive contexts. By training classifiers on neural patterns in response to one context and measuring their success in decoding patterns elicited in another, uncovering correspondence and abstraction across cognitive domains becomes possible (Kaplan et al. 2015). Cross-classification has been applied in various domains in the past, for example to investigate commonalities in visual perception and imagery (Shatek et al. 2019; Xie et al. 2020), between semantically similar words and images (Shinkareva et al. 2011), and language-invariant representations of objects (Quadflieg et al. 2011). Cross-classification across the conditions is usually conducted within-participant and decoding performance metrics are aggregated across the sample. Cross-subject decoding is a more stringent way of testing if neural patterns generalize and are shared across participants. In a previous study we have demonstrated that neural patterns in response to passive viewing of familiar and unfamiliar faces generalize across different groups of participants and familiarization conditions, providing evidence for a shared familiarity signal (Dalski et al. 2021).

For this present analysis, the two datasets were derived from two non-overlapping sets of participants, who were exposed to different sets of stimuli, different familiarization conditions, and performed two different tasks. Despite all these differences, we observed the emergence of a shared familiarity signal in the trials where both photographs depicted the perceptually familiarized identities (Sorted, Same). This finding suggests that cross-classification even between substantially different experimental tasks can be useful to gain insight into shared neural processes.

### Neural responses to perceptually and personally familiarized faces share common patterns, with a left hemispheric weight

Both time-resolved cross-classification and searchlight analyses yielded above-chance familiarity effects when trained on data from the Personal Familiarization experiment and tested on the matching task of the Perceptual Familiarization experiment. This suggests that seeing a familiarized face automatically and reliably elicits a familiarity response (Wiese et al. 2022). The earliest indication of above-chance classification was at 170 ms and was most pronounced over central-posterior sites. Interestingly, the present results show a left-hemisphere bias, in contrast to the notion of a general right-hemisphere dominance in face perception, and our own previous findings regarding the right hemispheric lateralization of identity and familiarity signals (Ambrus et al. 2019, 2021). Both familiarity and identity representations have been shown to encompass a wide spatial and temporal range. Where the spatial extent of these effects were investigated (e.g. Nemrodov et al. 2016; Dalski et al. 2021) it has been shown that identity or familiarity information can be read out from a large number of channels, with a right-lateralized occipito-temporal weight.

However, it has been suggested that both cerebral hemispheres are involved in the processing of face information, albeit in different ways. Studies indicate that the right hemisphere tends to take part in the more global, holistic processing of faces, while the left hemisphere performs more part- or feature-based processing (Rossion et al. 2000; Meng et al. 2012). For example, Gazzaniga and Smylie (1983) describe a complete callosal resection patient (V.P.) whose left hemisphere was not able to discriminate between faces but was able to make same-different decisions about face stimuli when the simple matches had to be made at the same point in the visual field. In accordance with these findings, we suggest that the enhanced effect in the left hemisphere might be due to the fact that the matching task required an active same-different decision, and for that, a deeper featural processing in the left hemisphere might have been necessary. We suggest that this might be one of the reasons the familiarity signal became prominent enough to allow a successful classification.

### No effect for unfamiliar and previously unseen identities

Interestingly, the only case where a consistent, significant familiarity signal was found was the familiarized, same-identity condition, i.e., when both simultaneously presented images were photographs of one of the two identities learned during the sorting task.

One explanation for this might be a related to the number of familiar identities presented simultaneously. While elevated responses were also seen in for one Sorted, one Foil identity (i.e., Sorted ID, Different) trials, these effects were far from robust. This could indicate that as participants alternated between attending to the two faces, looking at the Foil image attenuated the effects of familiarity. As no gaze-tracking was used in this study, future studies may help uncover the neural dynamics of such scenario.

Another important consideration is how “same” and “different” decisions are made. Mechanisms involved in deciding if two images depict “different” identities might more readily rely on image-based processing, instead of maintaining an image-independent representation.

Also, no consistent (un)familiarity effect was observed in the case of Unfamiliar (not sorted but seen in the decoding phase) identities and the previously Unseen identities. One might expect to see these trials to be cross-classified as ‘unfamiliar’ at a significant rate, instead, classification performance was observed to be at chance level for all conditions involving Unfamiliar and Unseen identities. It needs to be kept in mind that in the matching task, the participants were not informed about the experimental conditions, i.e., were not told that we would be showing images of identities they have seen in the sorting and decoding phases and were also not informed about the inclusion of the two not before seen identities. Furthermore, we also did not inform them about the fact that all Foil identities in the Different trials would be faces of novel, trial-unique identities. In this experiment no between-stimulus category match decisions had to be made (see **Figure 1**.), thus it is unknown whether some of the non-Sorted faces, including the Foils, were perceived as familiar based on superficial similarity, as care was taken to match the general appearance of the identities when we selected the stimuli (see **Figure 2**.). Because of this, while Sorted faces reliably elicited familiarity signals, neural responses to stimuli all other categories might have been much more varied, leading to near-chance classifier performance.

## Summary

In summary, we found evidence that face-learning through a brief perceptual card sorting task is sufficient to induce an image-independent abstraction of identity. This is illustrated by improved performance during the sorting task itself, and the lower reaction times and higher accuracies during a face matching task, for the familiarized compared to the unknown identities. The neural familiarity signal elicited by these identities during the matching task can be successfully cross-classified, when a classifier is trained on aggregated data from a separate set of participants passively viewing personally familiar faces. This implies that the visual processing of personally familiar faces and faces that were learned through perceptual exposure only, involve similar mechanisms, resulting in cross-decodable neural patterns. While the amount of semantic and socially relevant knowledge has been shown to increase the magnitude of familiarity effects in EEG, here we provided evidence that they are not essential for the emergence of this shared familiarity signal. Further studies are needed to uncover what processes contribute to this effect, and to evaluate what role additional knowledge plays in the generation of these neural patterns.

## Author contributions

AD, GK and GGA planned the study, GGA and AD conducted the analysis, AD, GGA and GK wrote the manuscript

